# The effect of unilateral hand muscle contraction on frontal alpha asymmetry and inhibitory control in intrinsic reward contexts, a randomized controlled trial

**DOI:** 10.1101/2024.09.05.611368

**Authors:** Atakan M. Akil, Renáta Cserjési, Tamás Nagy, Zsolt Demetrovics, H. N. Alexander Logemann

## Abstract

Challenged inhibitory control has been implicated in various disorders, including addiction. Previous research suggests that asymmetry of frontal brain activity, indexed by frontal alpha asymmetry (FAA), is associated with inhibitory control and could be a target for neuromodulatory intervention. Some evidence suggests that unilateral muscle contraction (UMC) can modulate FAA; however, experimental evidence is scarce. We conducted a randomized controlled trial, with 65 participants (M_age_ = 26.6; SD = 7.4), 37 of whom were females. We collected EEG data to calculate FAA and assessed inhibitory performance using the Stop Signal Task (SST) in neutral and intrinsic reward (palatable food) conditions, both before and after a unilateral left-hand muscle contraction task aimed at enhancing right relative to left frontal activity. We found a significant main effect of group on FAA. Specifically, UMC group was associated with higher right relative to left frontal activity, associated with resting state inhibitory activity. Event-related potential analyses revealed a significant dissociation between the stop N2 and stop P3 components as a function of time. More specifically, as time progressed, the stop N2 was enhanced, while the stop P3 was reduced. These results did not lead to observable changes in the behavioral index of stopping. In conclusion, UMC did not affect any behavioral and brain activity indices. There is some indication of a potential effect on FAA. However, this effect could reflect coincidental differences in trait FAA. Our findings provide new insights into the temporal dynamics of brain activity indices of inhibitory control.

## INTRODUCTION

Inhibitory control, the capacity to suppress a planned response, is a critical component of executive functions ^1^. Poor inhibitory control has been implicated in various psychiatric disorders such as substance abuse (e.g., cocaine ^2^, nicotine ^3^, and alcohol ^4^), obesity ^5^, major depressive disorder ^6^, and internet addiction ^7^. Studies suggest that the asymmetry of frontal brain activity is associated with inhibitory control. This asymmetry, presumably originating from the dorsolateral prefrontal cortex (DLPFC) and indexed by alpha oscillatory activity at the right frontal (F4) relative to the left frontal (F3) recording site ^8,9^. The presence of greater right frontal cortical activity compared to the left, as revealed by electroencephalogram (EEG) frontal alpha asymmetry, is believed to be associated with better inhibitory control ^9–11^. Several neuromodulatory methods to modulate frontal alpha asymmetry (FAA) have been extensively investigated, such as transcranial direct current stimulation ^12^, and EEG-neurofeedback ^13^. Specifically, there is evidence that brief manipulations of frontal asymmetry can plausibly lead to post-intervention effects in both clinical ^14^ and healthy sample group ^12, 13^.

Unilateral muscle contraction (UMC) has also been proposed as a possible technique to modulate FAA ^15^. This method involves the voluntary contraction of muscles (e.g., squeezing one hand) for a prolonged period. Previous studies found that UMC may have an effect on emotional and motivational tendencies, modulating the approach/avoidance system ^15^. The impact of UMC on one side of the body influencing emotional and motivational outcomes has been attributed to the activation of the contralateral hemisphere ^16^. Consistent with this assumption, it was also suggested that the effects caused by contractions were due to activation spreading to the opposite frontal regions ^17^. Specifically, inhibition-related traits and states have been associated with increased relative right frontal activity over the left ^11,15,18^. While UMC holds promise as a potential method for enhancing inhibitory control, no studies have thoroughly investigated the effect of UMC on inhibitory control, and its associated electrophysiological mechanism, especially within intrinsic reward contexts after the manipulation. Our current study addresses this gap.

To evaluate individual differences in inhibitory control, the Stop Signal Task (SST) has been widely employed ^19–21^. In this task, participants respond to “go” stimuli (e.g., palatable food pictures) with a key press, occasionally encountering a “stop” stimulus that requires withholding the response. Previous research has utilized a combination of SST and event-related potential (ERP) analyses to explore the neural mechanisms underlying inhibitory control ^22,23^. ERPs signify the synchronized activation of large neuron groups time-locked to an event ^24^. The Stop N2, manifesting around 200 ms latency, displays notably more negative amplitudes in successful stop trials compared to unsuccessful ones ^20^. Additionally, it has been suggested that the neurobiological correlate of the Stop N2 is the right inferior frontal gyrus. The Stop P3 is influenced by stopping success, exhibiting larger amplitudes for successful inhibitions than unsuccessful ones ^20^. Indeed, the Stop P3 is believed to signify inhibitory control ^25^ and is thought to originate from the superior frontal gyrus ^26^. Despite these presumed connections, the behavioral and electrophysiological indices of inhibitory control have not been thoroughly explored in the context of FAA and unilateral hand muscle contraction.

First, we aimed to enhance activity in the right relative to left frontal cortex which we think is indexed by the degree of asymmetry of frontal alpha synchronization based on previous research ^27^. Secondly, we employed a food reward condition, potentially activating left frontal cortical activity, to enhance the visibility of the intervention’s effect. The rewarding stimuli choice was inspired by prior research. Specifically, previous research has shown that healthy and normal-weight individuals naturally exhibit an implicit inclination towards high-calorie food ^28^. Similarly, another study investigated learned and intrinsic reward contexts, finding differences in inhibitory processes toward learned rewards (e.g., money) and intrinsic rewards (e.g., palatable food) in healthy individuals ^29^. We hypothesized that active contraction of left hand would lead to increased right relative to left frontal brain activity. This increase was expected to be manifested as a reduced eyes-open and eyes-closed FAA scores (F4-F3/F8-F7). Secondly, we expected that active contraction of left hand muscles would enhance inhibitory control, as measured by stop-signal reaction times, Stop N2, and Stop P3, in the reward condition compared to the neutral condition. Specifically, it would result in decreased reaction times and increased Stop N2 and Stop P3 amplitudes.

## METHODS

### Participants

A pilot study with a sample size of 10 participants validated our experimental procedure and formed the basis for determining the required sample size. We used G*Power ^30,31^ for a priori power analysis, using a power of 80%, a significance level of 5%, and assuming a test-retest correlation of 0.6 for the effect of time (before and after unilateral left-hand muscle contraction) on the primary outcome variable, FAA. The expected effect (f >0.237; η^2^partial > 0.053) was found to be reliably detected with a sample size of 30 individuals in the active unilateral left-hand muscle contraction group. We recruited a total of 65 individuals through social media and university courses. Thirty-seven (57%) of the participants were females and 61 (94%) of them were right-handed. Prior research has indicated that handedness does not have an impact on inhibitory control ^32^. Participants’ age ranged from 18 to 54, with an average of 26.6 (SD = 7.4). To be eligible for participation, individuals had to meet specific criteria. These criteria included being at least 18 years old and passing a screening for exclusion factors, such as the presence of psychological or psychiatric disorders, frequent headaches or migraines, epilepsy, significant prior head trauma, recent head injuries, chronic skin conditions, and current drug use. They were also asked to avoid smoking and drinking coffee at least two hours before the experiment. These criteria were assessed based on self-reported answers. We provided participants with a list of our exclusion criteria and asked if they met any of them. If a participant answered “yes” to any of the criteria, they were excluded from the study. Participants provided their demographic information and needed to self-assess their English proficiency using the Common European Framework of Reference for Languages - Self-assessment Grid ^33^ through Psytoolkit ^34,35^. All participants provided written informed consent. The research adhered to ethical guidelines outlined in the Declaration of Helsinki and its subsequent revisions. The research protocol received approval from the Institutional Review Board of Eötvös Loránd University. Each participant received a voucher or course credit as compensation for their involvement in the study.

### Stop Signal Task

The SST was programmed using OpenSesame ^36^ and was adapted from its original version ^20^, with the inclusion of a food reward condition based on insights from previous studies ^29,37^. Figure 1A illustrates the details of the task. Each experimental condition comprised a practice block with stop trials to determine the optimal go-stop interval for the subsequent first experimental block. A total of four experimental blocks were conducted for each condition, with each block consisting of 96 go trials and 32 stop trials, constituting 25 percent of the total trials. Therefore, each block comprised 128 trials, resulting in 512 trials per condition. Each study phase (pre-intervention and post-intervention) encompassed a total of 1024 trials, including both neutral and reward conditions. Consequently, the overall experimental design comprised 2048 trials, excluding practice blocks. Participants seated approximately 65 cm from the screen, received clear instructions both in written form and orally at the beginning of the task. The task started with a fixation dot in the center of the screen, holding participants’ attention for 2000 ms. In the reward condition, palatable food pictures (cookies, chips, chocolate, and nuts (115 px width (2.9°) x 200 px height (5.1°)) requiring a key response were presented for 150 ms, randomly in horizontal or vertical orientation. Conversely, in the neutral condition, the letters “X” or “O” (200 px width (5.1°) x 200 px height (5.1°)) necessitating a key response were displayed for 150 ms. Responses were executed with the left or right index fingers based on the stimulus. Stop trials, comprising infrequent presentation of the letter “S” (200 px width (5.1°) x 200 px height (5.1°)) following go stimuli, required participants to withhold their response. The Stimulus Onset Asynchrony was fixed at 350 ms, with the go-stop interval adjusted dynamically after each stop trial. If participants failed to inhibit their response, the go-stop interval was reduced by 50 ms, whereas it increased by 50 ms in case of successful inhibition. This tracking algorithm aimed for an approximate 50% inhibition rate, enhancing the reliability of stop-signal reaction time (SSRT) estimation ^21^, which is a well-established measure of inhibitory control derived from this task ^1,38^.

**Figure 1.**
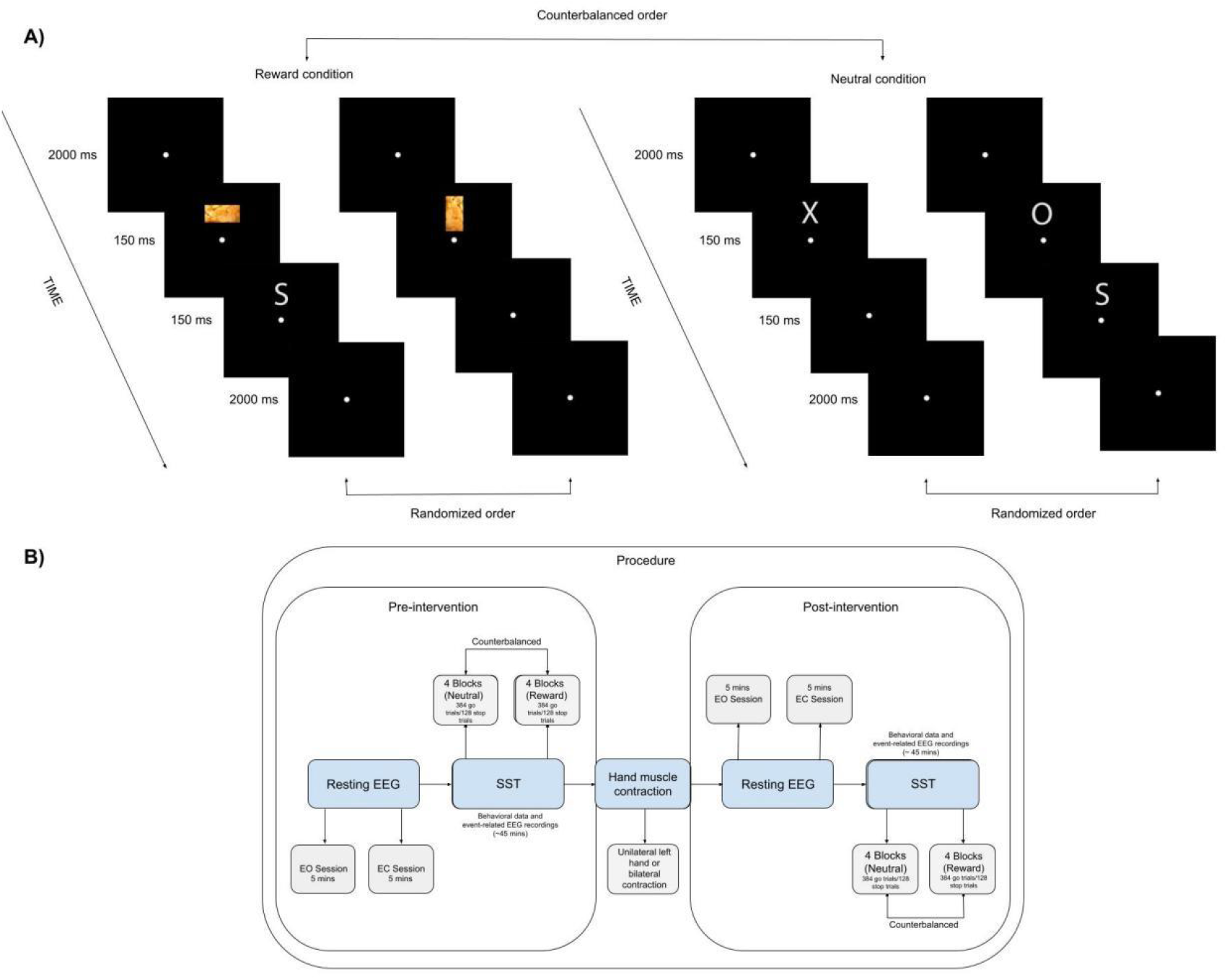
Figure 1A illustrates the procedure followed in the study. Participants began the Stop Signal Task in either the neutral or reward condition, based on a counterbalanced order. Each condition included a practice block with stop trials to determine the optimal go-stop interval for subsequent experimental blocks. Four experimental blocks were conducted per condition, each comprising 96 go trials and 32 stop trials, making up 25% of the total trials. This resulted in a total of 384 go trials and 128 stop trials per condition, totaling 768 go trials and 256 stop trials per phase (pre-intervention or post-intervention), including both neutral and reward conditions. The overall experimental design consisted of 1536 go trials and 512 stop trials. The two trials on the left depicts the reward condition, where target dimensions (horizontal/vertical) and palatable food pictures were randomly presented for 150 milliseconds. The final two trials on the right illustrates the neutral condition, where the letters “X” and “O” were randomly presented for 150 milliseconds. Some trials were followed by the letter “S,” indicating stop trials where participants were required to withhold their initiated responses to go stimuli. In Figure 1B, the process began with gathering pre-intervention resting-state EEG data to measure frontal alpha asymmetry, conducted in two sessions: one with eyes open for 5 minutes and another with eyes closed for 5 minutes. Subsequently, participants performed the Stop Signal Task in a counterbalanced order of neutral and reward conditions. Following this, either bilateral hand muscle contraction or unilateral lefthand muscle contraction was administered for 10 minutes, with continuous 45-second contraction trials followed by 15-second rest intervals. Frontal alpha asymmetry was recorded during the muscle contraction as well. After the intervention, the same procedure was repeated for post-modulation assessment, including resting-state EEG and the Stop Signal Task.

### Electrophysiological Data Collection

Scalp voltage measurements were obtained using a 21-channel EEG cap with Ag/AgCl electrodes, following the 10-20 system. We used Nexus-32 from Mind Media ^39^. Electrophysiological data was collected using EEG with the hardware common average reference. Vertical electrooculography signals were recorded above and below the left eye, while horizontal electrooculography signals were captured by electrodes placed at the outer canthi of both eyes. EEG was continuously recorded, including during hand muscle contraction. The data was offline re-referenced to linked mastoids. Subsequently, the data was filtered and down sampled to 512 Hz.

### Frontal Alpha Asymmetry

FAA scores were derived from EEG data collected across three distinct conditions. Initially, we computed these scores from two separate 5-minute resting-state sessions, one with eyes open (EO) and another with eyes closed (EC), both conducted before and after the intervention. Gathering EEG resting-state recordings in these two scenarios enhances our understanding of the impacts of sensory input, internal cognitive processes, and the inherent activity of the brain. Employing this methodology contributes to a comprehensive comprehension of the functional organization of the brain in different states ^40,41^. Additionally, FAA scores were calculated based on hand muscle contractions. The preprocessing steps were performed using BrainVision Analyzer 2 (www.brainproducts.com), following established procedures ^**42**^. Initially, signal preprocessing involved applying a high-pass filter at 0.5 Hz, a low-pass filter at 40 Hz, and a 50 Hz notch filter. Subsequently, the first and final 10 seconds of EEG data were excluded due to the presence of artifacts. The remaining data was segmented into 2-second epochs. For the EO condition, ocular artifacts were corrected using independent component analysis based on data from vertical electrooculography and horizontal electrooculography electrodes. Any epochs still containing artifacts, as determined by exceeding a criterion of ±75 µV maximum amplitude relative to the baseline, were excluded from analysis. To analyze spectral content, Power Spectral Density was computed using Fast Fourier Transform with a 10% Hanning window on the remaining epochs after whole-segment baseline correction. Subsequently, these epochs were averaged, and the mean activity in the alpha frequency range (8-13 Hz) was computed. The resulting values for relevant electrodes were exported for further analysis. Using R, alpha power values were adjusted for skewness through natural log transformation ^42^. Finally, frontal alpha asymmetry was calculated by subtracting the log-transformed alpha values at lateral left electrode sites from those at right electrode sites, specifically F4-F3 and F8-F7.

### Event-related Potentials

Using BrainVision Analyzer 2, first, the signals were referenced to the linked mastoid electrodes. Subsequently, following previous studies ^43,44^, the EEG data underwent offline filtering with a high-pass cutoff of 0.5 Hz, a low-pass cutoff of 30 Hz, and a 50

Hz notch filter. The data was then divided into epochs, ranging from -100 ms to 2600 ms. To address ocular artifacts, we applied independent component analysis. Following this correction, the epochs were segmented and baseline-corrected using a reference window of -100 to 0 ms. We performed a segmentation aligned with the presentation of the go-signal, followed by baseline correction and artifact rejection. Artifact rejection criteria included amplitudes exceeding ±75 µV, with a 200 ms buffer before and after events marked as bad. Subsequently, we conducted a segmentation aligned with the stop-signal onset and applied baseline correction. Separate averages were computed for segments corresponding to unsuccessful stop trials and successful stop trials. The inhibition-related ERPs were derived by subtracting the average activity for unsuccessful stop trials from that of successful stop trials. After carefully examining the grand average waveforms, specific latency intervals were identified for further analysis, data for the stop N2 component at F4 electrode were extracted from the time window of 172-292 ms, while data for the stop P3 component at Cz electrode were selected from the 191-241 ms time window for export. As an exploratory analysis, we also analyzed the stop N2 component from the 160-180 ms time window.

### Unilateral Muscle Contraction

The primary objective of implementing UMC in this study was to enhance the relative brain activity in the right DLPFC, a region potentially linked with inhibitory control. In the experimental group, participants engaged in a UMC protocol wherein they squeezed a ball using their left hands for 45-second followed by 15-second rest intervals. The UMC task lasted for 10 minutes. Meanwhile, the control group involved simultaneous ball squeezing with both hands ^15^.

### Procedure

Figure 1B shows the detailed procedure. This research employed a controlled design, with both within-subject (time before and after the intervention/condition being neutral and reward) and between-subject (unilateral or bilateral hand muscle contraction) factors. Upon arriving at the laboratory, participants read an information letter, confirmed their eligibility based on the inclusion criteria, and completed the informed consent form. Following this, EEG electrodes were placed on their scalp for data collection during resting-state sessions. Resting-state EEG data were collected for a total of 10 minutes, divided into two sessions of five minutes each, one with eyes open and the other with eyes closed. After the EEG recording, participants filled out questionnaires. Next, they completed the initial phase of the Stop Signal Task before undergoing the unilateral muscle contraction intervention. Participants were then assigned to receive either unilateral or bilateral hand muscle contraction in a counterbalanced order. During the post-intervention assessment, the same sequence of steps, resting-state EEG and the Stop Signal Task, was repeated. Both the intervention and pre-/post-assessments were conducted on the same day and took approximately five hours.

### Statistical Analysis

We conducted data analyses using R ^45^. Following the computation of key variables, we excluded participants with missing data and outliers exceeding 3 standard deviations (SD) in case of erroneous data. Subsequently, we performed a repeated measures ANOVA with a 2×2 design to investigate the effect of UMC on the primary outcome, FAA (EO/EC). For the potential electrophysiological (Stop N2/Stop P3) and behavioral index (SSRT) of inhibitory control, we performed a repeated measures of ANOVA with a 2×2x2 design. We also conducted an exploratory analysis for the effect of the interventions on the first SST condition (neutral/reward separately). We provided Bayes factors (BF) of 10 (BF_10_) for the results. We used “BayesFactor” package in R. The results of exploratory and Bayesian factor analyses can be found in the supplementary materials.

## RESULTS

### Resting-state frontal alpha asymmetry

To assess the impact of UMC on resting-state FAA scores, we conducted a repeated measures ANOVA. This analysis included time (pre- and post-intervention) as the factor within subjects and group (unilateral or bilateral hand muscle contraction) as the factor between subjects. Table 1 shows the details. The findings from this analysis indicated that the main effect of group on FAA F8-F7 EO was significant: F(1, 110) = 7.14, *p* = 0.008, η_p_^2^ < 0.061. This result was also supported by Bayesian statistics, BF_10_ = 4.94 (Supplementary Table 1). Specifically, it was observed that UMC group was associated with higher inhibitory motivation regardless of temporal influence (Figure 2). The observed effect probably imply inherent disparities between the groups, which could arise from a variety of factors such as initial group characteristics. On the other hand, the immediate (online) effect of unilateral lefthand muscle contraction was not found to be statistically significant F(1, 59) = 0.07, *p* = 0.787, η_p_^2^ = 0.001. We also conducted an analysis of the time x group interaction by incorporating three time points (pre-, during, and post-intervention) for FAA. The effects were found to be insignificant. The results is provided in the supplementary materials (Supplementary Table 2).

**Table 1.** Results of the frontal alpha asymmetry models.

| Models | Df | F | P | $\eta_p^2$ |
| --- | --- | --- | --- | --- |
| **FAA F4-F3 (EO) (*n* = 57)** |  |  |  |  |
| Time | 1 | 0.01 | 0.952 | <0.001 |
| Group | 1 | 0.06 | 0.796 | <0.001 |
| Time x Group | 1 | 0.20 | 0.651 | 0.001 |
| Residuals | 110 |  |  |  |
| **FAA F4-F3 (EC) (*n* = 58)** |  |  |  |  |
| Time | 1 | <0.01 | 0.982 | <0.001 |
| Group | 1 | 0.39 | 0.532 | 0.003 |
| Time x Group | 1 | <0.01 | 0.936 | <0.001 |
| Residuals | 112 |  |  |  |
| **FAA F4-F3 (int) (*n* = 61)** |  |  |  |  |
| Group | 1 | 0.07 | 0.787 | 0.001 |
| Residuals | 59 |  |  |  |
| **FAA F8-F7 (EO) (*n* = 57)** |  |  |  |  |
| Time | 1 | <0.01 | 0.926 | <0.001 |
| Group | 1 | 7.14 | 0.008* | 0.061 |
| Time x Group | 1 | 0.02 | 0.871 | <0.001 |
| Residuals | 110 |  |  |  |
| **FAA F8-F7 (EC) (*n* = 58)** |  |  |  |  |
| Time | 1 | 0.42 | 0.516 | 0.003 |
| Group | 1 | 0.39 | 0.532 | 0.003 |
| Time x Group | 1 | 0.52 | 0.471 | 0.004 |
| Residuals | 112 |  |  |  |
| **FAA F8-F7 (int) (*n* = 61)** |  |  |  |  |
| Group | 1 | 0.01 | 0.921 | <0.001 |
| Residuals | 59 |  |  |  |

Note: Participants with missing values were excluded from the analyses. "Int" refers to the frontal alpha asymmetry scores during the bilateral and unilateral hand muscle contraction interventions.

**Figure 2.**
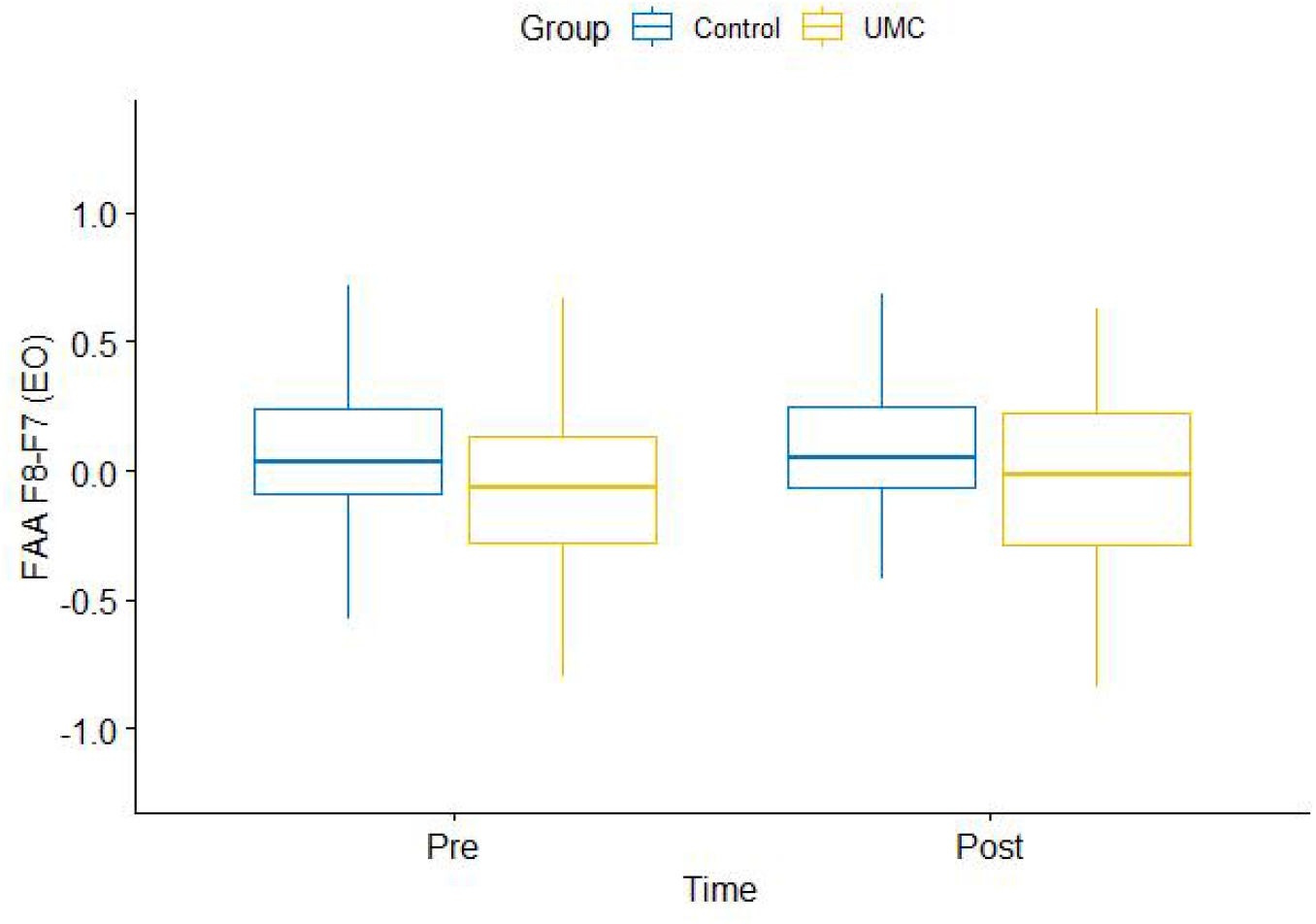
The figure shows the main effect of group on eyes open frontal alpha asymmetry (F8-F7 electrodes).

### Behavioral indices of inhibitory control

To analyze the impact of UMC on inhibitory control, we conducted a repeated measures ANOVA with time (pre- and post-intervention) and condition (neutral and reward) as the within-subject factors, and group (unilateral and bilateral hand muscle contraction) as the between-subject factor. First, we excluded the participants with missing values and <10% inhibition rates for the calculation of stop-signal reaction times. The details can be found in Table 2 and 3. Notably, we did not observe any significant influence of UMC on SSRT scores, specifically: F(1, 220) = 0.21, *p* = 0.645, η_p_^2^ < 0.001. Bayesian factor results can be found in Supplementary Table 3.

**Table 2.**
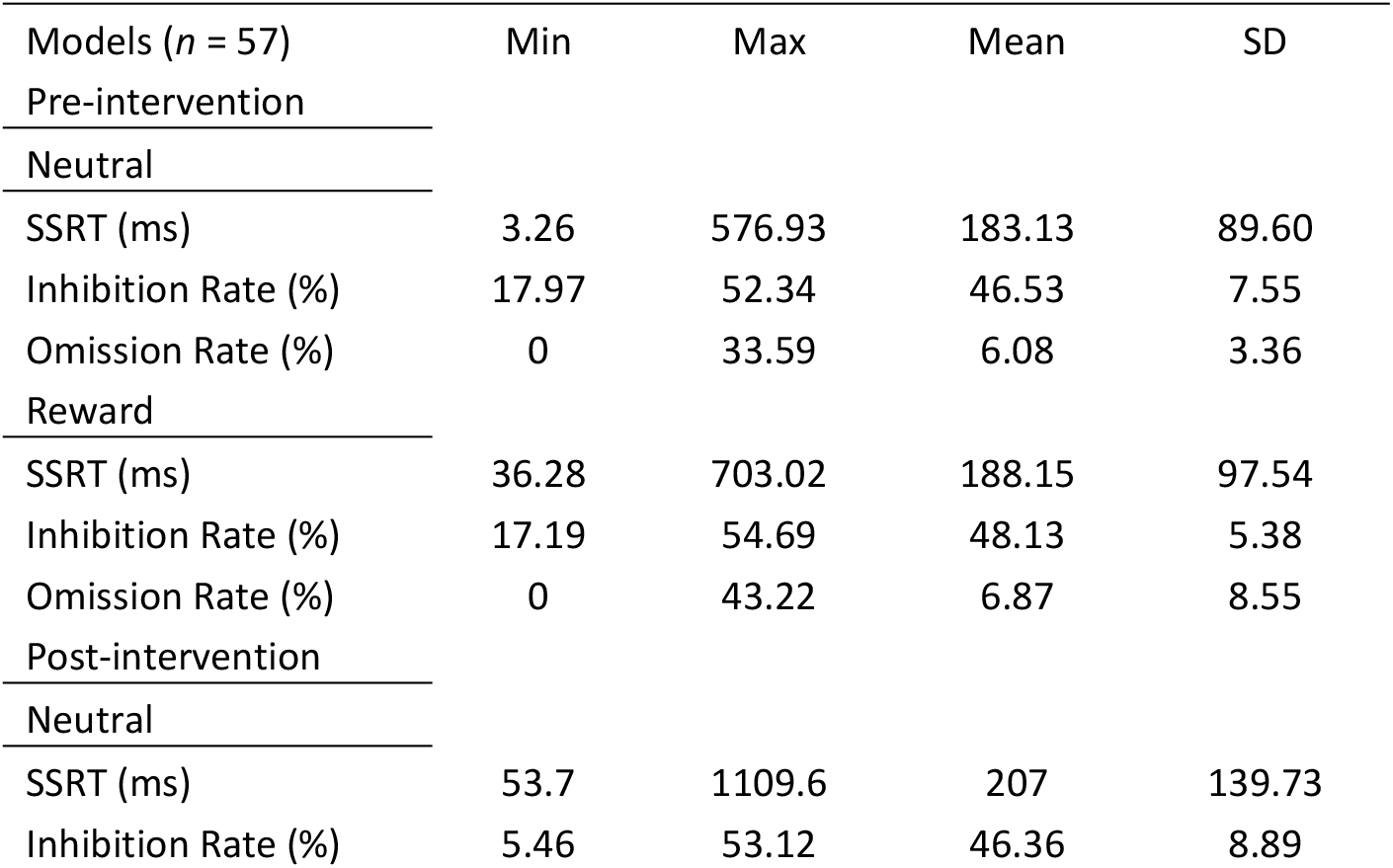
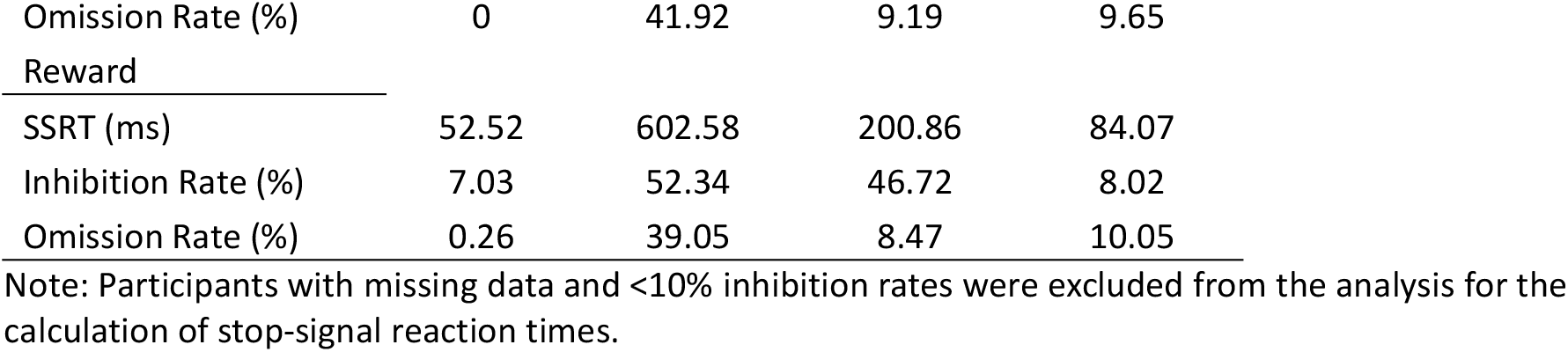
Descriptive statistics regarding the stop-signal task performance.

**Table 3.**
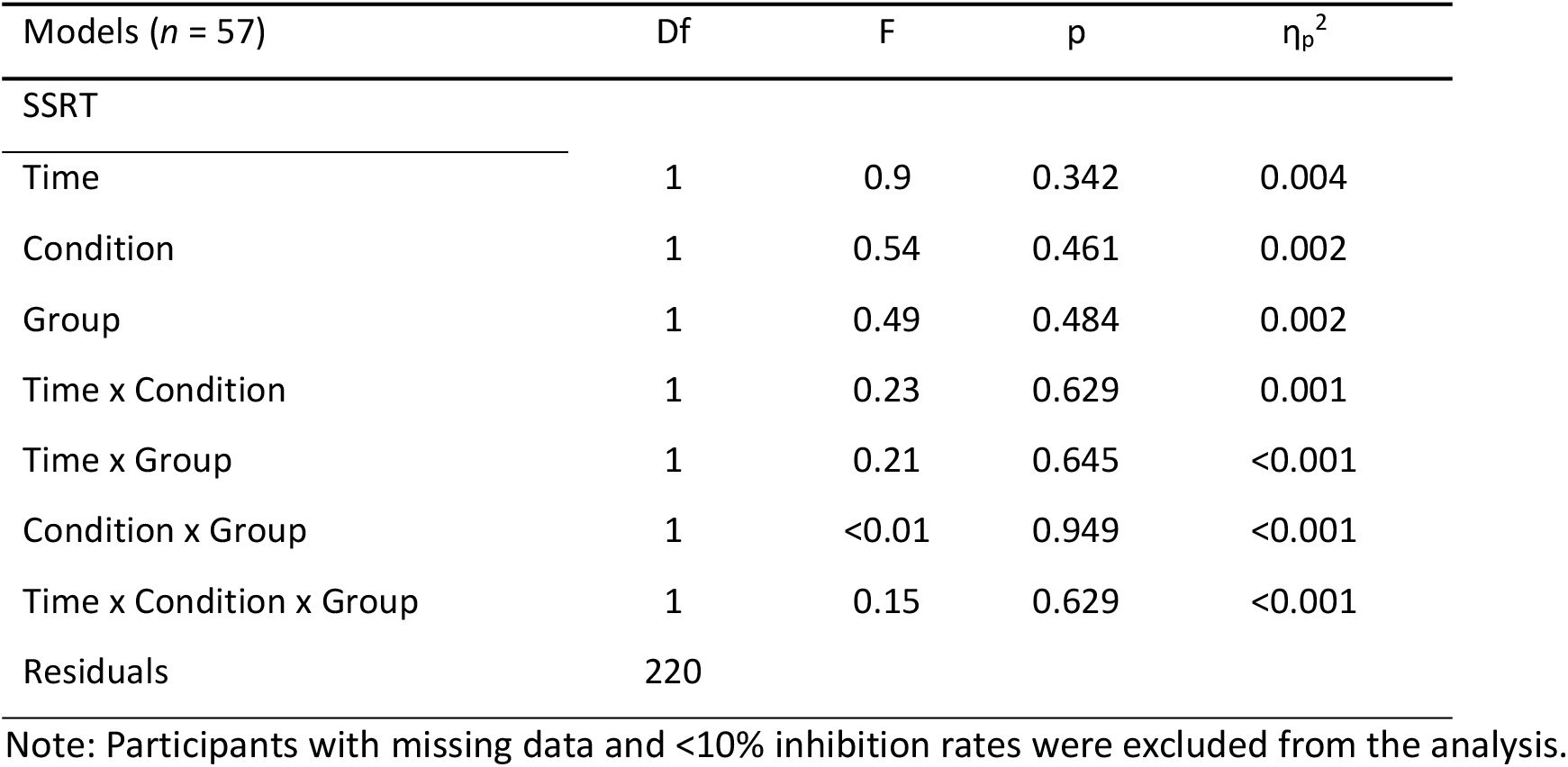
Results of the stop-signal reaction times models.

### Brain activity indices of inhibitory control

To explore whether UMC affects brain activity related to inhibitory control, we conducted a repeated measures ANOVA. This analysis consisted of time (pre- and post-intervention) and condition (neutral or reward) as the within-subject factors, while group (unilateral and bilateral hand muscle contraction) was the between-subject factor. Table 4 shows the statistical details. There was no significant effect of unilateral hand muscle contraction on the brain activity indices of inhibitory control. However, the main effect of time on the stop N2 and the stop P3 was statistically significant: F(1, 204) = 4.79, *p* = 0.029, η_p_^2^ = 0.022, BF_10_ = 1.37 in Supplementary Table 4 and F(1, 196) = 4.19, *p* = 0.041, η_p_^2^ = 0.020, BF_10_ = 1.10 in Supplementary Table 5, respectively. Figure 3 and 4 show the results of the Stop N2 and the Stop P3, respectively. Surprisingly, while the stop N2 suggested an increased negativity over time, the stop P3 showed a decreased positivity activity. Our exploratory analysis regarding the stop N2 at 160-180 ms showed a similar effect of time: F(1, 204) = 4.40, *p* = 0.037, η_p_^2^ = 0.021. For details, please refer to Supplementary Tables 6 and 7.

**Table 4.**
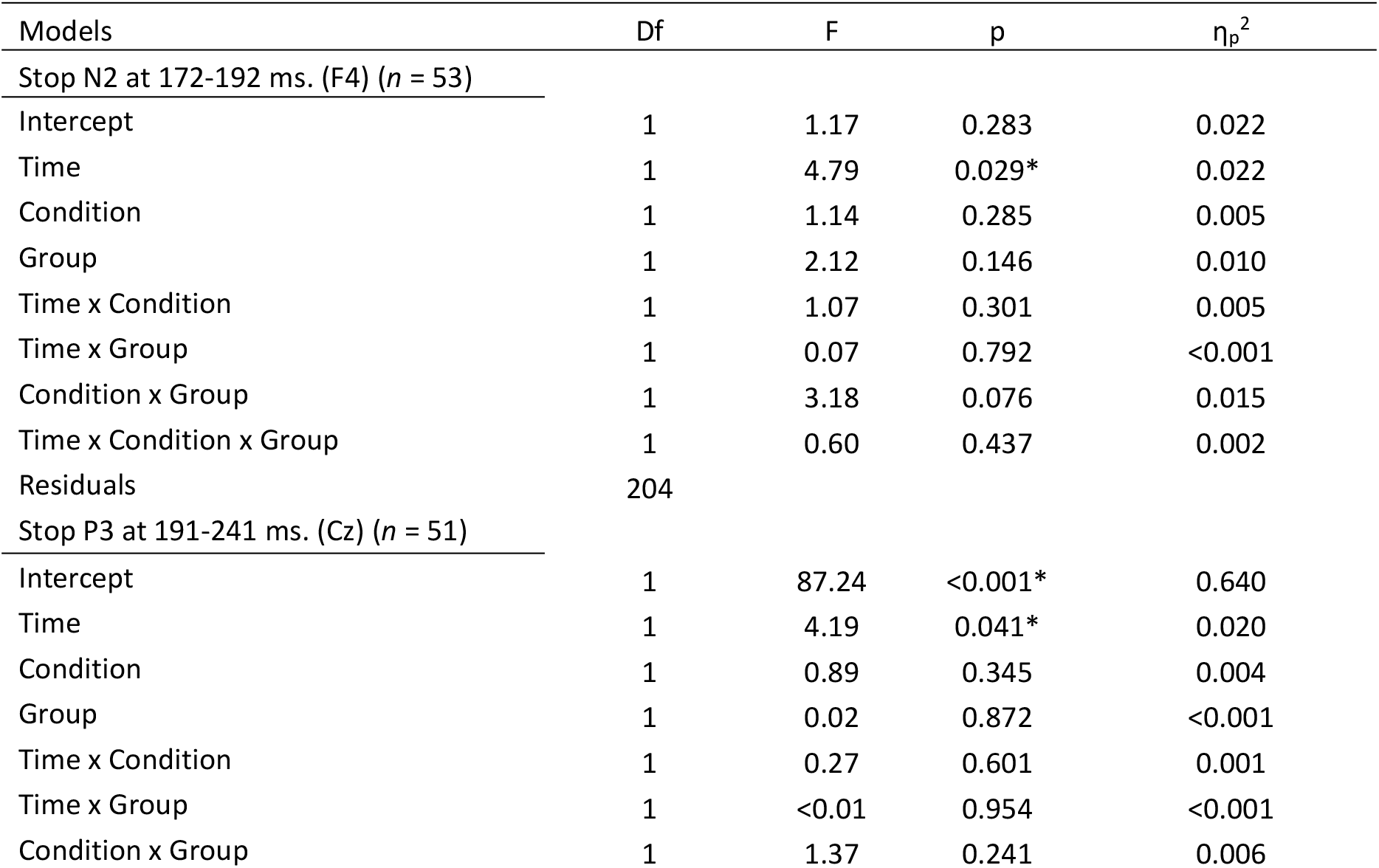
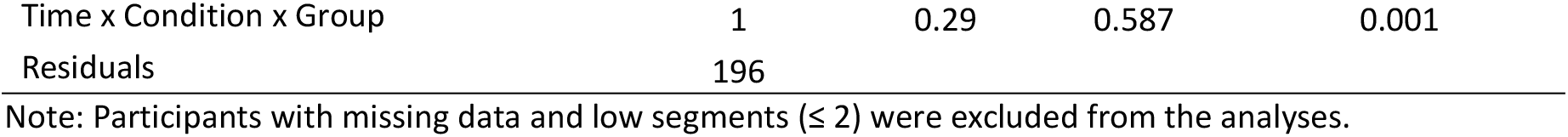
Results of the event-related potential models.

| Models | Df | F | p | $\eta_p^2$ |
| --- | --- | --- | --- | --- |
| **Stop N2 at 172-192 ms. (F4) (*n* = 53)** |  |  |  |  |
| Intercept | 1 | 1.17 | 0.283 | 0.022 |
| Time | 1 | 4.79 | 0.029* | 0.022 |
| Condition | 1 | 1.14 | 0.285 | 0.005 |
| Group | 1 | 2.12 | 0.146 | 0.010 |
| Time x Condition | 1 | 1.07 | 0.301 | 0.005 |
| Time x Group | 1 | 0.07 | 0.792 | <0.001 |
| Condition x Group | 1 | 3.18 | 0.076 | 0.015 |
| Time x Condition x Group | 1 | 0.60 | 0.437 | 0.002 |
| Residuals | 204 |  |  |  |
| **Stop P3 at 191-241 ms. (Cz) (*n* = 51)** |  |  |  |  |
| Intercept | 1 | 87.24 | <0.001* | 0.640 |
| Time | 1 | 4.19 | 0.041* | 0.020 |
| Condition | 1 | 0.89 | 0.345 | 0.004 |
| Group | 1 | 0.02 | 0.872 | <0.001 |
| Time x Condition | 1 | 0.27 | 0.601 | 0.001 |
| Time x Group | 1 | <0.01 | 0.954 | <0.001 |
| Condition x Group | 1 | 1.37 | 0.241 | 0.006 |
| Time x Condition x Group | 1 | 0.29 | 0.587 | 0.001 |
| Residuals | 196 |  |  |  |

Note: Participants with missing data and low segments ( $\leq 2$ ) were excluded from the analyses.

**Figure 3.**
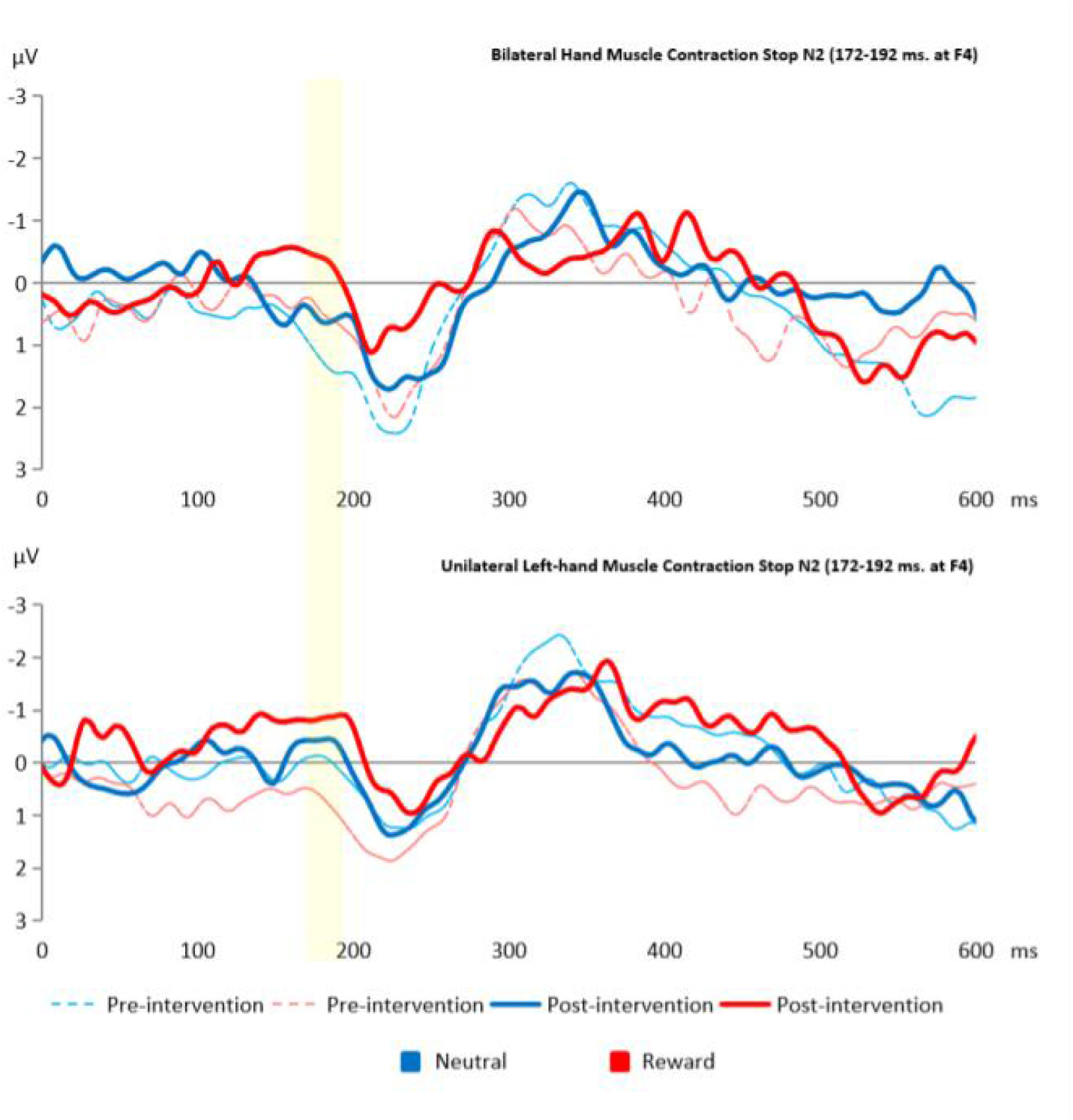
This figure illustrates the effect of time on the Stop N2 (172-192 ms) at F4 electrode. The bar represents the Stop N2 peaks.

**Figure 4.**
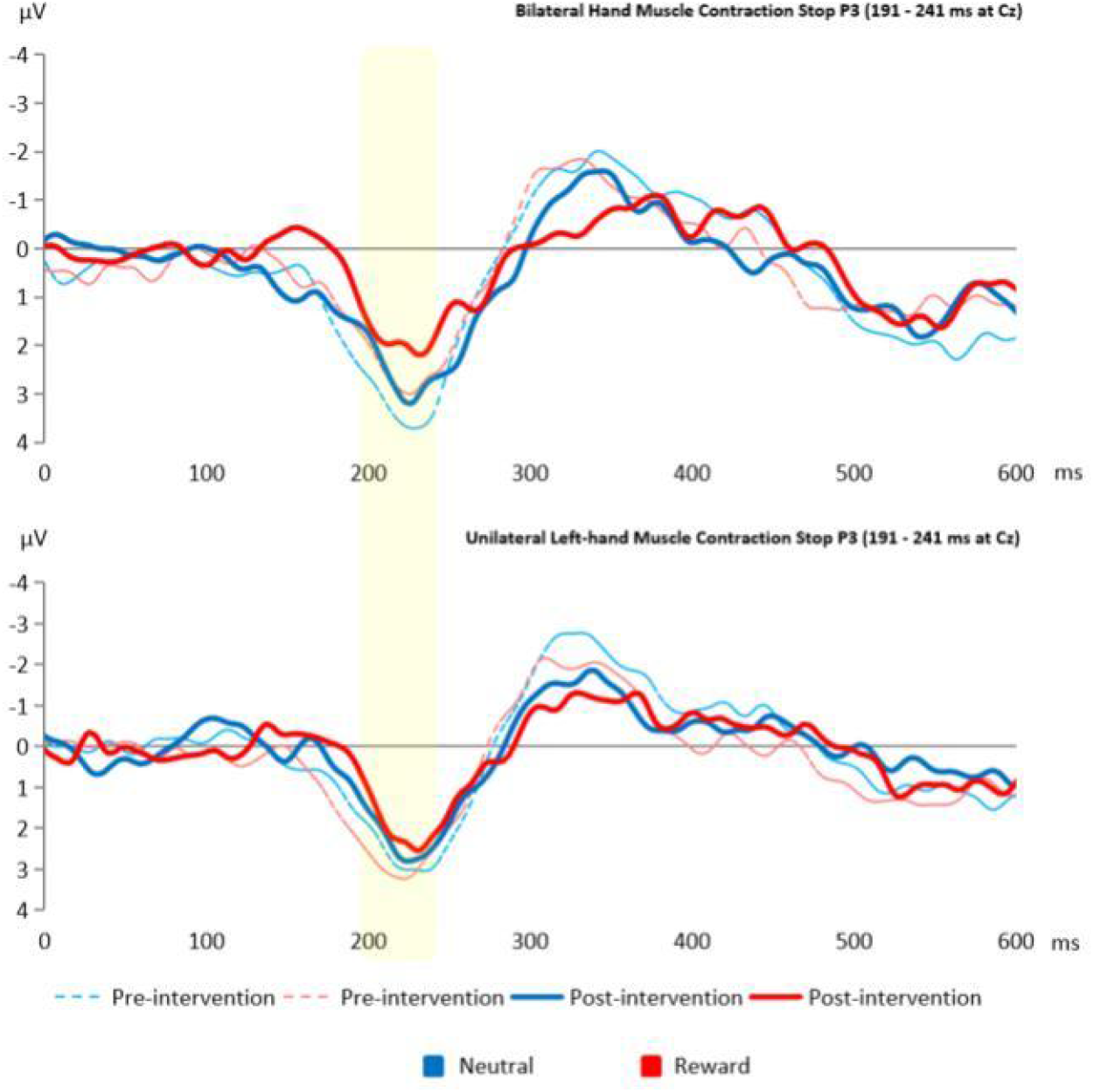
This figure illustrates the effect of time on the Stop P3 (191-241) at Cz electrode. The bar represents the Stop P3 peaks.

### Further exploratory analyses

We conducted further analyses regarding the effect of unilateral left-hand muscle contraction on SSRTs. We created two models, one of which included participants starting with the neutral condition, while the other consisted of participants starting with the reward condition. Therefore, we divided the task to investigate the immediate effects of the interventions. We found no effect of UMC on the first condition, both neutral and reward (F(1, 28) = 2.07, *p* = 0.161, η_p_^2^ = 0.068 and F(1, 26) = 0.06, *p* = 0.809, η_p_^2^ = 0.002, respectively). These findings were also supported by Bayesian factors. Supplementary Tables 8, 9, and 10 contain detailed results. Lastly, we performed an exploratory analysis with the participants who had less than 20 percent omission rates (*n* = 45) and found that there was no significant effect of time x condition x group interaction: F(1, 172) = 1.67, *p* = 0.198, η_p_^2^ = (Supplementary Table 11).

## DISCUSSION

This study explored the impact of unilateral muscle contraction on frontal alpha asymmetry, a potential marker of inhibitory control, and behavioral and neural indices associated with inhibitory control. We found that the main effect of group on frontal alpha asymmetry was significant. Interestingly, a noteworthy dissociation emerged in event-related potential components related to inhibition. More specifically, temporal dynamics emerged as a significant factor influencing neural markers of inhibitory control. Our results showed that over the course of time, the stop N2 exhibited a significant decrease, signifying a higher negativity and suggesting heightened inhibitory neural activity. However, the stop P3 shows a significant decrease in positivity, suggesting less inhibitory neural activity. These effects did not lead to observable behavioral changes.

As hypothesized, there was an association between UMC and frontal alpha asymmetry. More specifically, the UMC group was associated with higher right relative to left frontal cortical activity, compared to the bilateral hand muscle contraction group. This result holds some promise regarding a potential effect of unilateral muscle contraction and is in line with previous studies ^15,16^,^17^. However, the time x group interaction was not significant. This means that there was no significant group difference in the change from pre-to post-intervention. Hence we cannot rule out that the aforementioned effect is due to a coincidental group difference in trait FAA at baseline.

The main effect of time on the stop N2 indicates heightened inhibitory brain activity, while the stop P3 suggests a decline in inhibitory brain activity over the same temporal span. In the current paradigm, we employed a dynamic tracking algorithm which resulted in variation in go-stop intervals. Hence, the stop-signal associated electrophysiological response was computed for different go-stop intervals, which partly mitigates the contribution of go-associated response activity . However, it has been shown that early latency components such as the N1 modulation by stopping success to auditory tones can be difficult to detect without filtering out response-associated activity from stop-signal associated activity (Bekker et al., 2005). Importantly, the stop N2 and stop P3 were both identified and were significant in a previous report using the same experimental paradigm (Akil et al., 2024). Nevertheless, we cannot exclude the possibility that response-associated activity contributes to the observed N2 and P3 modulations.

The debate regarding the functional relevance of the stop N2 and the stop P3 in response conflict, task outcome expectancy, and inhibitory control is still ongoing, though most acknowledge that both brain activity components are key in inhibition processes. For instance, Groom & Cragg ^46^ investigated the N2 and P3 event-related potentials as markers of response conflict and response inhibition. The N2 amplitude was identified as a marker of response conflict, showing greater amplitude on incongruent trials. Conversely, the P3 amplitude was highlighted as a marker of response inhibition, with enhanced amplitude on trials requiring response inhibition. Hence, our study may indicate that the heightened negativity amplitude implies a rise in response conflict over time. This could be attributed to a potential decrease in inhibitory control caused by increasing tiredness and fatigue, as evident in the stop P3.

The changes in the stop P3 could also be influenced by participants’ evolving expectations about task outcomes. Specifically, previous studies found that the P3 magnitude increases with the subjective unexpectedness of task outcomes ^47,48^. This suggests that, over time, individuals exhibit less anticipation or expectation of a specific outcome when engaging in the SST. Therefore, our results align with the idea that the neural processes underlying inhibitory control evolve with experience and temporal dynamics.

The control condition may not be entirely passive and could modulate brain activity, given the absence of differences between groups, but the effect over time. Hence, it may be important to consider the potential role of unilateral and bilateral hand muscle contractions, similar to the underlying effects of non-invasive brain modulation techniques like the effects observed in transcranial direct current stimulation ^49–51^. Changes in participants’ motivation or engagement with the task across sessions may impact inhibitory control processes differently at different stages of information processing, as reflected in the stop N2 and stop P3 components. Therefore, refining the parameters that modulate the stop N2 and the stop P3 could enhance the assessment of inhibitory control in healthy and clinical populations, contributing to a more nuanced understanding of the phenomenon, and leading to establishment of specific treatment targets.

## Conclusion

We investigated the relationship between bilateral/unilateral hand muscle contraction, frontal alpha asymmetry, and inhibitory control in food reward contexts. We did not find a significant effect of UMC on indices of inhibitory control. The main effect of time on the stop N2 and the stop P3 was significant. The finding sheds light on the nuanced temporal modulation of inhibitory brain activities, enhancing our understanding of the association between temporal dynamics and neural markers of response conflict, post-task evaluation, and inhibitory control.

## Data Availability Statement

Raw data can be found:

Self-report data:

https://cloud.adaptingminds.nl/d/s/xZdt5NlOfbLJq3NgkhGyDfrC5DNQNyKH/CL3VG7CGDnIXYcdeQzdTqyO1EDDuxGuF-RLKgC9goMQs

Computer task data:

https://cloud.adaptingminds.nl/d/s/pNSCmsKwgtyIcpuPG1p0ZXJpELft5lHQ/-A0i5HSKjfUnOA5v-KtDRMg5GDaVJQo4-8rEgNXooMQs

EEG data:

https://cloud.adaptingminds.nl/d/s/pNSCsFioxnEa7u33HfEbCGZ6rc6zQ6BE/n4MoIDaoeJjPoaGTT3w0eWLxCRVXSSdc-OLIgqM0oMQs

