## Supplementary Materials for "The effect of unilateral hand muscle contraction on frontal alpha asymmetry and inhibitory control in intrinsic reward contexts, a randomized controlled trial"

**Supplementary Table 1. Bayesian repeated measures ANOVA results for the frontal alpha asymmetry models**

| Models | $BF_{10}$ | Error (%) |
|---|---|---|
| FAA F4-F3 (EO) (*n* = 57) | | |
| Group | 0.20 | ±0.03 |
| Time | 0.19 | ±0.03 |
| Time + Group | 0.03 | ±1.76 |
| Time + Group + Time x Group | 0.01 | ±1.45 |
| FAA F4-F3 (EC) (*n* = 58) | | |
| Group | 0.23 | ±0.03 |
| Time | 0.19 | ±0.03 |
| Time + Group | 0.04 | ±2.15 |
| Time + Group + Time x Group | 0.01 | ±2.05 |
| FAA F4-F3 (int) (*n* = 61) | | |
| Group | 0.26 | ±0.01 |
| FAA F8-F7 (EO) (*n* = 57) | | |
| Group | 4.94 | ±0.01 |
| Time | 0.19 | ±0.03 |
| Time + Group | 0.95 | ±2.07 |
| Time + Group + Time x Group | 0.25 | ±2 |
| FAA F8-F7 (EC) (*n* = 58) | | |
| Group | 0.23 | ±0.03 |
| Time | 0.23 | ±0.03 |
| Time + Group | 0.05 | ±2.14 |
| Time + Group + Time x Group | 0.01 | ±2.03 |
| FAA F8-F7 (int) (*n* = 61) | | |
| Group | 0.26 | ±0.01 |

Note: Participants with missing values were excluded from the analyses. "Int" refers to the frontal alpha asymmetry scores during the bilateral and unilateral hand muscle contraction interventions.

**Supplementary Table 2. The effect of time x group interaction on frontal alpha asymmetry**

| Models | Df | F | P | $\eta_p^2$ |
|---|---|---|---|---|
| FAA F4-F3 (EO) (*n* = 56) | | | | |
| Time | 1 | 0.41 | 0.661 | 0.005 |
| Group | 1 | 0.02 | 0.882 | <0.001 |
| Time x Group | 1 | 0.12 | 0.887 | 0.001 |
| Residuals | 162 | | | |
| FAA F8-F7 (EO) (*n* = 56) | | | | |
| Time | 1 | 0.02 | 0.972 | <0.001 |
| Group | 1 | 5.34 | 0.022* | 0.031 |
| Time x Group | 1 | 1.17 | 0.310 | 0.014 |
| Residuals | 162 | | | |

Note: Participants with missing values were excluded from the analyses. The analysis includes all three time points (pre-intervention, during the intervention, and post-intervention).

**Supplementary Table 3. Bayesian repeated measures ANOVA results for the stop-signal reaction time models**

| Models (*n* = 57) | $BF_{10}$ | Error (%) |
|---|---|---|
| Group | 0.18 | ±0.07 |
| Time | 0.22 | ±0.06 |
| Group + Time | 0.03 | ±1.55 |
| Time + Group + Time x Group | <0.01 | ±2.5 |
| Condition | 0.18 | ±0.07 |
| Group + Condition | 0.03 | ±2.02 |
| Time + Condition | 0.04 | ±1.12 |
| Group + Time + Condition | <0.01 | ±2.78 |
| Group + Time + Group x Time + Condition | <0.01 | ±9.86 |
| Group + Condition + Group x Condition | <0.01 | ±2.56 |
| Group + Time + Condition + Group x Condition | <0.01 | ±2.04 |
| Group + Time + Group x Time + Condition + Group x Condition | <0.01 | ±2.46 |
| Time + Condition + Time x Condition | <0.01 | ±2.4 |
| Group + Time + Condition + Time x Condition | <0.01 | ±3.15 |
| Group + Time + Group x Time + Condition + Time x Condition | <0.01 | ±3.85 |
| Group + Time + Condition + Group x Condition + Time x Condition | <0.01 | ±3.03 |
| Group + Time + Group x Time + Condition + Group x Condition + Time x Condition | 7.37 | ±3.14 |
| Group + Time + Group x Time + Condition + Group x Condition + Time x Condition + Group x Time x Condition | 1.91 | ±4.74 |

Note: Participants with missing data and <10% inhibition rates were excluded from the analysis for the calculation of stop-signal reaction times.

**Supplementary Table 4. Bayesian repeated measures ANOVA results for the Stop N2 (172-192 ms) models**

| Models (*n* = 53) | $BF_{10}$ | Error (%) |
|---|---|---|
| Group | 0.39 | ±0.04 |
| Time | 1.37 | ±0.01 |
| Group + Time | 0.53 | ±1.75 |
| Time + Group + Time x Group | 0.11 | ±1.38 |
| Condition | 0.25 | ±0.05 |
| Group + Condition | 0.09 | ±1.11 |
| Time + Condition | 0.34 | ±2.57 |
| Group + Time + Condition | 0.14 | ±1.39 |
| Group + Time + Group x Time + Condition | 0.03 | ±1.88 |
| Group + Condition + Group x Condition | 0.09 | ±10.09 |
| Group + Time + Condition + Group x Condition | 0.12 | ±4.01 |
| Group + Time + Group x Time + Condition + Group x Condition | 0.02 | ±3.95 |
| Time + Condition + Time x Condition | 0.10 | ±2.12 |
| Group + Time + Condition + Time x Condition | 0.04 | ±5.46 |
| Group + Time + Group x Time + Condition + Time x Condition | <0.01 | ±3.59 |
| Group + Time + Condition + Group x Condition + Time x Condition | 0.03 | ±2.84 |
| Group + Time + Group x Time + Condition + Group x Condition + Time x Condition | <0.01 | ±6.1 |
| Group + Time + Group x Time + Condition + Group x Condition + Time x Condition + Group x Time x Condition | <0.01 | ±4.66 |

Note: Participants with missing data and low segments (≤ 2) were excluded from the analyses.

**Supplementary Table 5. Bayesian repeated measures ANOVA results for the Stop P3 (191-141 ms) models**

| Models (*n* = 51) | $BF_{10}$ | Error (%) |
|---|---|---|
| Group | 0.15 | ±0.07 |
| Time | 1.10 | ±0.02 |
| Group + Time | 0.16 | ±2.14 |
| Time + Group + Time x Group | 0.03 | ±2.06 |
| Condition | 0.23 | ±0.05 |
| Group + Condition | 0.03 | ±1.77 |
| Time + Condition | 0.25 | ±3.29 |
| Group + Time + Condition | 0.04 | ±10.41 |
| Group + Time + Group x Time + Condition | <0.01 | ±3.5 |
| Group + Condition + Group x Condition | 0.01 | ±1.93 |
| Group + Time + Condition + Group x Condition | 0.01 | ±8.45 |
| Group + Time + Group x Time + Condition + Group x Condition | <0.01 | ±9.88 |
| Time + Condition + Time x Condition | 0.06 | ±1.41 |
| Group + Time + Condition + Time x Condition | <0.01 | ±2.87 |
| Group + Time + Group x Time + Condition + Time x Condition | <0.01 | ±2.32 |
| Group + Time + Condition + Group x Condition + Time x Condition | <0.01 | ±3.66 |
| Group + Time + Group x Time + Condition + Group x Condition + Time x Condition | <0.01 | ±4 |
| Group + Time + Group x Time + Condition + Group x Condition + Time x Condition + Group x Time x Condition | <0.01 | ±4.46 |

Note: Participants with missing data and low segments (≤ 2) were excluded from the analyses.

**Supplementary Table 6. Results of the Stop N2 (160-180 ms) models**

| Models | Df | F | p | $\eta_p^2$ |
|---|---|---|---|---|
| Stop N2 at 160-180 ms. (F4) (*n* = 53) | | | | |
| Time | 1 | 4.40 | 0.037* | 0.021 |
| Condition | 1 | 1.39 | 0.239 | 0.006 |
| Group | 1 | 1.12 | 0.289 | 0.005 |
| Time x Condition | 1 | 1.43 | 0.233 | 0.006 |
| Time x Group | 1 | 0.04 | 0.841 | <0.001 |
| Condition x Group | 1 | 2.86 | 0.092 | 0.013 |
| Time x Condition x Group | 1 | 0.12 | 0.722 | <0.001 |
| Residuals | 204 | | | |

Note: Participants with missing data and low segments (≤ 2) were excluded from the analyses.

**Supplementary Table 7. Bayesian repeated measures ANOVA results for the Stop N2 (160-180 ms) models**

| Models (*n* = 53) | $BF_{10}$ | Error (%) |
|---|---|---|
| Group | 0.25 | ±0.05 |
| Time | 1.15 | ±0.02 |
| Group + Time | 0.28 | ±2.14 |
| Time + Group + Time x Group | 0.05 | ±2.05 |
| Condition | 0.28 | ±0.05 |
| Group + Condition | 0.06 | ±1.76 |
| Time + Condition | 0.33 | ±3.28 |
| Group + Time + Condition | 0.08 | ±10.35 |
| Group + Time + Group x Time + Condition | 0.01 | ±3.49 |
| Group + Condition + Group x Condition | 0.04 | ±1.92 |
| Group + Time + Condition + Group x Condition | 0.06 | ±8.44 |
| Group + Time + Group x Time + Condition + Group x Condition | 0.01 | ±9.79 |
| Time + Condition + Time x Condition | 0.12 | ±1.36 |
| Group + Time + Condition + Time x Condition | 0.03 | ±2.84 |
| Group + Time + Group x Time + Condition + Time x Condition | <0.01 | ±2.24 |
| Group + Time + Condition + Group x Condition + Time x Condition | 0.02 | ±3.64 |
| Group + Time + Group x Time + Condition + Group x Condition + Time x Condition | <0.01 | ±2.87 |
| Group + Time + Group x Time + Condition + Group x Condition + Time x Condition | <0.01 | ±3.12 |

+ Group x Time x Condition

**Supplementary Table 8. Results of the effect of interventions on stop-signal reaction times among participants receiving the neutral condition first**

| Models (*n* = 16) | Df | F | p | $\eta_p^2$ |
|---|---|---|---|---|
| SSRT | | | | |
| Time | 1 | 0.20 | 0.653 | 0.007 |
| Group | 1 | 0.15 | 0.697 | 0.005 |
| Time x Group | 1 | 2.07 | 0.161 | 0.068 |
| Residuals | 28 | | | |

Note: Participants with missing data and <10% inhibition rates were excluded from the analysis for the calculation of stop-signal reaction times. Only participants receiving the neutral condition after the intervention were included in the analysis.

**Supplementary Table 9. Results of the effect of interventions on stop-signal reaction times among participants receiving the reward condition first**

| Models (*n* = 15) | Df | F | p | $\eta_p^2$ |
|---|---|---|---|---|
| SSRT | | | | |
| Time | 1 | <0.01 | 0.981 | <0.001 |
| Group | 1 | <0.01 | 0.940 | <0.001 |
| Time x Group | 1 | 0.06 | 0.809 | 0.002 |
| Residuals | 26 | | | |

Note: Participants with missing data and <10% inhibition rates were excluded from the analysis for the calculation of stop-signal reaction times. Only participants receiving the reward condition after the intervention were included in the model.

**Supplementary Table 10. Bayesian results regarding the effect of interventions on stop-signal reaction times in first conditions**

| Models | $BF_{10}$ | Error (%) |
|---|---|---|
| Neutral condition first (*n* = 16) | | |
| Group | 0.35 | ±0.0 |
| Time | 0.36 | ±0.0 |
| Time + Group | 0.12 | ±2.53 |
| Time + Group + Time x Group | 0.11 | ±1.11 |
| Reward condition first (*n* = 15) | | |
| Group | 0.34 | ±0 |
| Time | 0.34 | ±0 |
| Time + Group | 0.11 | ±2.47 |
| Time + Group + Time x Group | 0.05 | ±2.01 |

**Supplementary Table 11. Results of the stop-signal reaction times models**

| Models (*n* = 45) | Df | F | p | $\eta_p^2$ |
|---|---|---|---|---|
| SSRT | | | | |
| Time | 1 | 2.63 | 0.107 | 0.015 |
| Condition | 1 | 0.25 | 0.612 | 0.001 |
| Group | 1 | <0.01 | 0.985 | <0.001 |
| Time x Condition | 1 | 0.11 | 0.733 | <0.001 |
| Time x Group | 1 | 0.78 | 0.378 | 0.004 |
| Condition x Group | 1 | 0.19 | 0.660 | 0.001 |
| Time x Condition x Group | 1 | 1.67 | 0.198 | 0.009 |
| Residuals | 172 | | | |

Note: Participants with missing data and <10% inhibition rates, and over 20 % omission rates were excluded from the analysis for the calculation of stop-signal reaction times.
